# The Neurotoxin DSP-4 Dysregulates the Locus Coeruleus-Norepinephrine System and Recapitulates Molecular and Behavioral Aspects of Prodromal Neurodegenerative Disease

**DOI:** 10.1101/2022.09.27.509797

**Authors:** Alexa F. Iannitelli, Michael A. Kelberman, Daniel J. Lustberg, Anu Korukonda, Katharine E. McCann, Bernard Mulvey, Arielle Segal, L. Cameron Liles, Steven A. Sloan, Joseph D. Dougherty, David Weinshenker

**Affiliations:** Department of Human Genetics, Emory University School of Medicine, Atlanta, GA 30322, USA; Department of Genetics, Washington University School of Medicine, St. Louis, MO 63110, USA; Department of Psychiatry, Washington University School of Medicine, St. Louis, MO 63110, USA

**Author notes:** Address correspondence to: David Weinshenker, PhD, Department of Human Genetics, Emory University School of Medicine, 615 Michael St., Whitehead 301.

## Abstract

The noradrenergic locus coeruleus (LC) is among the earliest sites of tau and alpha-synuclein pathology in Alzheimer’s disease (AD) and Parkinson’s disease (PD), respectively. The onset of these pathologies coincides with loss of noradrenergic fibers in LC target regions and the emergence of prodromal symptoms including sleep disturbances and anxiety. Paradoxically, these prodromal symptoms are indicative of a noradrenergic hyperactivity phenotype, rather than the predicted loss of norepinephrine (NE) transmission following LC damage, suggesting the engagement of complex compensatory mechanisms. Because current therapeutic efforts are targeting early disease, interest in the LC has grown, and it is critical to identify the links between pathology and dysfunction. We employed the LC-specific neurotoxin DSP-4, which preferentially damages LC axons, to model early changes in the LC-NE system pertinent to AD and PD in male and female mice. DSP-4 (2 doses of 50 mg/kg, 1 week apart) induced LC axon degeneration, triggered neuroinflammation and oxidative stress, and reduced tissue NE levels. There was no LC cell death or changes to LC firing, but transcriptomics revealed reduced expression of genes that define noradrenergic identity and other changes relevant to neurodegenerative disease. Despite the dramatic loss of LC fibers, NE turnover and signaling were elevated in terminal regions and were associated with anxiogenic phenotypes in multiple behavioral tests. These results represent a comprehensive analysis of how the LC-NE system responds to axon/terminal damage reminiscent of early AD and PD at the molecular, cellular, systems, and behavioral levels, and provides potential mechanisms underlying prodromal neuropsychiatric symptoms.

## Introduction

Alzheimer’s disease (AD) and Parkinson’s disease (PD) are the most common cognitive and motor neurodegenerative disorders, respectively. Both conditions are characterized by abnormal protein accumulation in neurons leading to cellular dysfunction and death. While these disorders differ in their etiology and clinical presentation, early pathology in the brainstem locus coeruleus (LC) is a hallmark of both AD and PD (Weinshenker, 2018). The LC is the primary source of central norepinephrine (NE) and projects to nearly every other brain region (Foote et al., 1983; Berridge and Foote, 1996; Aston-Jones et al., 1999; Poe et al., 2020). LC neurons are the first to accumulate hyperphosphorylated tau in AD (Braak et al., 2011) and they develop aberrant alpha-synuclein before dopamine neurons of the substantia nigra (SN) in PD (Del Tredici et al., 2002). Although the LC eventually undergoes catastrophic degeneration in both diseases, these neurons can harbor pathology for years before cell death, displaying axon and dendrite loss in initial stages of AD and PD (Halliday et al., 1990; Busch et al., 1997; Theofilas et al., 2017; Doppler et al., 2021; Gilvesy et al., 2022). Combined, these data suggest that LC-NE deficiency contributes to AD and PD. Indeed, experimental lesions of the LC exacerbate neurodegeneration and cognitive deficits in rodent models, and loss of LC integrity correlates with cognitive decline in humans (Weinshenker, 2018; Jacobs et al., 2021b). However, this simplistic view is inconsistent with other data indicating excessive noradrenergic transmission, particularly early in disease. For example, increased levels and turnover of NE have been reported in the cerebrospinal fluid of AD patients (Palmer et al., 1987; Hoogendijk et al., 1999; Henjum et al., 2022). Moreover, because LC activity promotes arousal and stress responses, increased, rather than decreased, NE signaling is consistent with many of the prodromal symptoms of AD and PD including anxiety, depression, agitation, and sleep disturbances (Weinshenker, 2018).

Animal models of AD (Goodman et al., 2021; Kelly et al., 2021; Kelberman et al., 2022) and PD (Butkovich, 2019; Matschke et al., 2022) that recapitulate early LC pathology but lack outright noradrenergic cell death exhibit LC-NE hyperactivity, anxiety-like behavior, and hyperarousal, which can be alleviated with the administration of adrenergic antagonists. Likewise, neuropsychiatric symptoms in AD correlate with high LC signal contrast and respond to blockade of adrenergic receptors (Peskind et al., 2005; Cassidy et al., 2022). We have proposed a more complex model in which damaged LC neurons engage compensatory mechanisms that lead to noradrenergic hyperactivity and contribute to prodromal behavioral phenotypes, followed later by frank LC cell death and NE deficiency that accelerates cognitive decline (Weinshenker, 2018). However, causal relationships between LC damage, cellular and molecular compensatory mechanisms, and prodromal symptoms remain to be investigated and established.

To better understand noradrenergic dysfunction in early neurodegenerative disease and its links to compensatory mechanisms and behavioral abnormalities, we employed the LC-specific neurotoxin N-(2-chloroethyl)-N-ethyl-2-bromobenzylamine (DSP-4), which preferentially damages noradrenergic axons compared to cell bodies (Grzanna et al., 1989; Zhang et al., 1995). While many groups have reported depletion of NE following DSP-4 administration (Grzanna et al., 1989; Theron et al., 1993; Wolfman et al., 1994; Harro et al., 1999; Szot et al., 2010), it is important to acknowledge some limitations: (1) the effects of DSP-4 are often interpreted as noradrenergic ablation without taking potential compensatory mechanisms into account; (2) most have focused on only a single (or a few) aspect of LC function (e.g. NE abundance *or* axon integrity *or* LC-sensitive behaviors); and (3) many different dosing regimens and species have been used, limiting our ability to integrate the findings into a comprehensive picture of how the LC-NE system responds to damage. Here, we assessed the consequences of DSP-4 administration on molecular, cellular, and behavioral responses of the LC-NE system in parallel. Our results are critical for understanding LC dysfunction in AD and PD, and may provide a foundation for early diagnostic and intervention strategies for these disorders.

## Materials and Methods

### Animals

Adult male and female C57BL/6 mice were used for all behavioral, electrophysiological, and immunohistochemical experiments. For translating ribosome affinity purification (TRAP) RNA-sequencing experiments, we used male and female transgenic *Slc6a2-eGFP/Rpl10a* mice (B6;FVB-*Tg(Slc6a2-eGFP/Rpl10a)JD1538Htz/J*, The Jackson Laboratory, #031151), which incorporate an EGFP/Rpl10a ribosomal fusion protein into a bacterial artificial chromosome under the *Slc6a2* (NE transporter; NET) promoter to allow for the isolation of polysomes and translating mRNAs specifically from noradrenergic neurons. *Slc6a2-eGFP/Rpl10a* mice were purchased and maintained as hemizygotes on a C57BL/6 background. Mice were group housed with sex- and age-matched conspecifics (maximum of 5 animals per cage) until one week prior to behavioral testing, and then individually housed for the subsequent week of experimentation until sacrifice. Animals were maintained on a 12:12 light:dark cycle (lights on at 0700), and food and water were available *ad libitum,* unless otherwise specified. All experiments were conducted at Emory University in accordance with the National Institutes of Health *Guideline for the Care and Use of Laboratory Animals* and approved by the Emory Institutional Animal Care and Use Committee. Mice were treated with DSP-4 (50 mg/kg, i.p.; Sigma-Aldrich, St. Louis, MO) or vehicle (0.9% NaCl) on days 1 and 7. Electrophysiology and TRAP were conducted on day 14, and behavioral testing commenced on day 14 and ended on day 18.

### HPLC

Mice were anesthetized with isoflurane and euthanized by rapid decapitation. The pons, prefrontal cortex, and hippocampus were rapidly dissected on ice and flash-frozen in isopentane (2-Methylbutane) on dry ice. The samples were weighed and stored at −80°C until processing for HPLC. As previously described (Lustberg et al., 2022), tissue was thawed on ice and sonicated in 0.1 N perchloric acid (10 μl/mg tissue) for 12 s with 0.5 s pulses. Sonicated samples were centrifuged (16,100 rcf) for 30 min at 4 °C, and the supernatant was then centrifuged through 0.45 μm filters at 4000 rcf for 10 min at 4 °C. For HPLC, an ESA 5600A CoulArray detection system, equipped with an ESA Model 584 pump and an ESA 542 refrigerated autosampler was used. Separations were performed using an MD-150 × 3.2 mm C18, 3 μm column (Thermo Scientific) at 30 °C. The mobile phase consisted of 8% acetonitrile, 75 mM NaH2PO4, 1.7 mM 1-octanesulfonic acid sodium and 0.025% trimethylamine at pH 2.9. A 20 μL of sample was injected. The samples were eluted isocratically at 0.4 mL/min and detected using a 6210 electrochemical cell (ESA, Bedford, MA) equipped with 5020 guard cell. Guard cell potential was set at 475 mV, while analytical cell potentials were −175, 100, 350 and 425 mV. The analytes were identified by the matching criteria of retention time measures to known standards (Sigma Chemical Co., St. Louis MO). Compounds were quantified by comparing peak areas to those of standards on the dominant sensor.

### Immunohistochemistry

Mice were euthanized with an overdose of sodium pentobarbital (Fatal Plus, 150 mg/kg, i.p.; Med-Vet International, Mettawa, IL) and were transcardially perfused with cold 4% PFA in 0.01 M PBS. After extraction, brains were post-fixed overnight in 4% PFA at 4°C and then transferred to a 30% sucrose/PBS solution for 72 h at 4°C. Brains were embedded in OCT medium (Tissue-Tek) and sectioned by cryostat into 40-um-thick coronal sections at the level of the LC, ACC, and hippocampus. Sections were blocked in 5% normal goat serum (NGS) in 0.01 M PBS/0.1% Triton-X permeabilization buffer and then incubated for 24 h at 4°C in NGS blocking buffer with primary antibodies listed in Table 1. Following washes in 0.01 M PBS, sections were incubated for 2 h in blocking buffer, including secondary antibodies described in Table 1. After washing, sections were mounted onto Superfrost Plus slides and coverslipped with Fluoromount-G plus DAPI (Southern Biotech, Birmingham, AL).

**Table 1:**
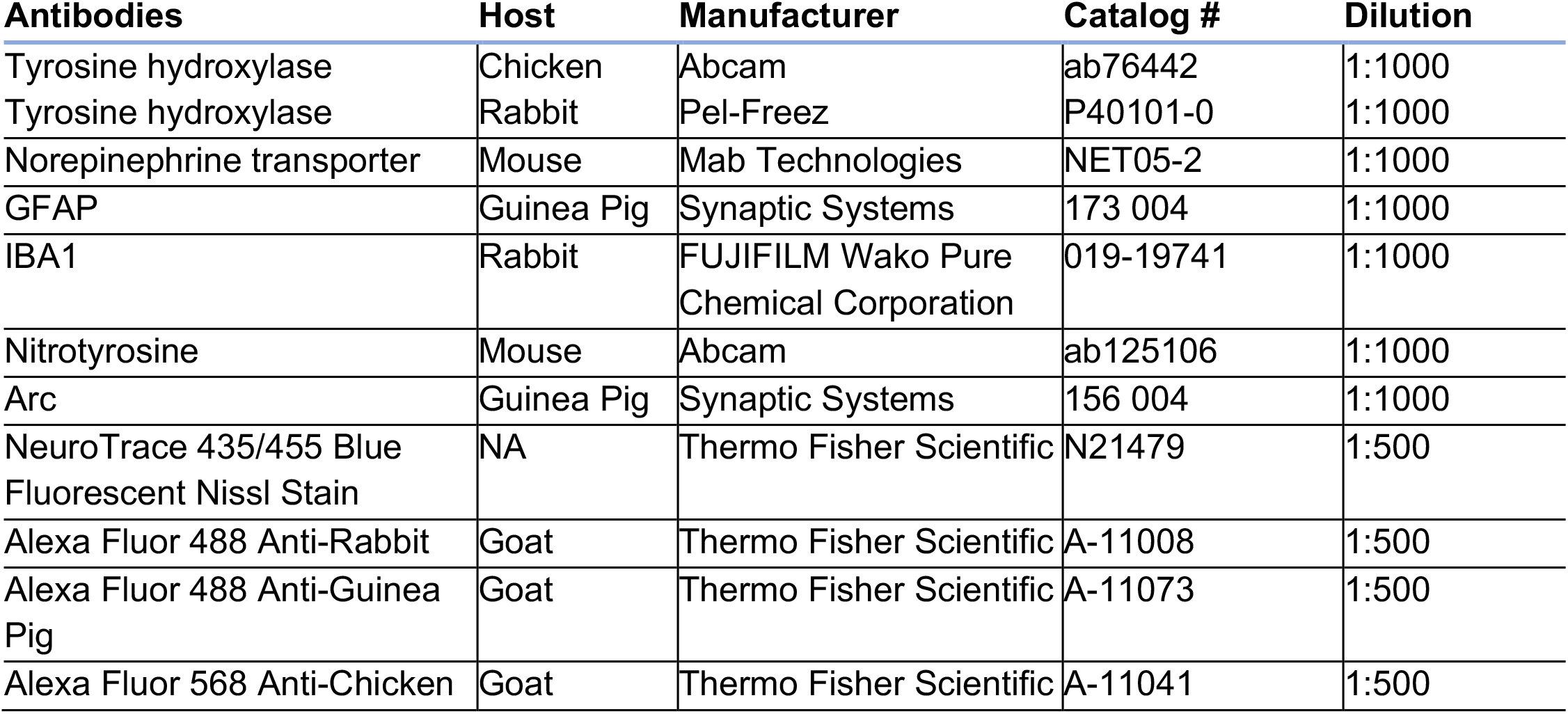
IHC antibodies.

### Quantification

For the catecholaminergic markers NE transporter (NET) and tyrosine hydroxylase (TH), the activity marker activity-regulated cytoskeletal gene (Arc), and the oxidative stress marker 3-nitrotyrosine (3-NT), immunofluorescent micrographs were acquired on a Leica DM6000B epifluorescent upright microscope at 20x magnification with uniform exposure parameters for each stain and region imaged. For glial markers, immunofluorescent images were acquired as z-stack images (10 z-stacks; pitch: 0.1 μm) at 20x magnification and compressed on a Keyence BZ-X700 microscope system. One representative atlas-matched section was selected from each animal and a standard region of interest was drawn for each image to delineate the LC, ACC, and hippocampus. For catecholaminergic and glial markers, image processing and analysis were conducted using the FIJI/ImageJ software. The analysis pipeline included standardized background subtraction, intensity thresholding (Otsu method), and pixel intensity measurements within defined ROIs of the same size (Lustberg et al., 2020a). Furthermore, *GFAP^+^ IBA1^+^* and *S100B* ^+^ cells were quantified based on size and shape for glia (50-1000 μm^2^, circularity 0.15-1.0) (Rorabaugh et al., 2017).

### Silver staining

Assessment of degenerating neuronal processes and cell bodies was performed using the NeuroSilver Staining Kit II (FD NeuroTechnologies, Inc., Columbia, MD) on fixed, free-floating sections from saline and DSP-4 treated mice (n = 3-4 per treatment group). Brain sections (40 um) were collected by cryostat (Leica) at the level of the LC, ACC, and DG. Staining was performed according to the manufacturer’s instructions (Kolisnyk et al., 2017; Kumbhare et al., 2017), after which tissue sections were allowed to air-dry overnight on SuperFrost Plus slides (Fisher Scientific, Hampton, NH). Once dry, sections were cleared for 2 min in CitriSolv xylene substitute (Fisher Scientific) and coverslipped with DPX non-aqueous mounting media (MilliporeSigma, Burlington, MA). Brightfield micrographs of silver-stained sections were acquired at 20x magnification using a Keyence BZ-X700 at 20x magnification with uniform light exposure parameters throughout image acquisition.

### Cell counts

Coronal tissue sections were processed as described above (*Immunohistochemistry*) and images were acquired as described for glial markers (*Quantification*). Using TH as a guide for anatomical LC borders and DAPI as a marker for individual nuclei, sections were atlas-matched and quantified using HALO imaging software (Indica Labs, v3.3.2541.420, FISH/IF v2.1.4). Nine sections were analyzed for each animal (n=3/group), covering most of the LC. The total area analyzed did not differ between the saline and DSP-4 treated groups. Within HALO, nuclei were defined using DAPI, and TH-positive and Nissl-positive nuclei cells were counted and summed across all nine sections for each animal. Comparisons were made between the number of Nissl+ and TH+ nuclei cells in the saline and DSP-4 treated groups.

### Translating Ribosome Affinity Purification (TRAP)

To obtain adequate quantities of RNA for sequencing, samples from two 6-8 month-old, same-sex and treatment *Slc6a2-eGFP/Rpl10a* mice were pooled to form a biological replicate by dissecting out the hindbrain posterior to the pontine/hypothalamic junction (cerebellum was discarded). Six biological replicates were collected per treatment group. Each replicate was homogenized and TRAP was performed as described (Mulvey et al., 2018), resulting in LC-enriched “TRAP” samples and whole-hindbrain “input” samples. RNA was extracted using Zymo RNA Clean & Concentrator-5 kit, and subsequently sent for library preparation and Illumina sequencing by NovoGene to a minimum depth of 20 million fragments per sample. Forward and reverse sequencing files from each replicate were aligned to the mouse genome (mm10) using STAR alignment, and counts were obtained using FeatureCounts in R Bioconductor. Two samples from saline-treated mice were removed from analysis because the TRAP protocol failed to enrich *Slc6a2* above a 10-fold change, a quality control threshold observed in all other saline-treated samples. All subsequent analysis utilized R Bioconductor packages. Sequencing data will be available on NCBI GEO at the time of publication.

To further characterize the gene expression changes, we performed a Weighted Gene Coexpression Network Analysis (WGCNA), as described previously (Zhang and Horvath, 2005; Langfelder et al., 2008). All samples that survived quality control parameters were used to create the co-expression network. Default parameters were primarily used throughout the analysis. The soft threshold power was set at 10, the point in which the scale free topology fit index was above 0.80. Minimum module size was set to 30 genes and modules with >95% similarity were merged, resulting in 159 modules which were then correlated with treatment. We also compared gene expression patterns in our dataset with a repository of gene sets using Gene Set Enrichment Analysis (GSEA 4.2.3) (Mootha et al., 2003; Subramanian et al., 2005). Gene set permutation was used, as recommended by GSEA documentation for experiments with fewer than 7 samples per group. Other parameters were set to default settings using 1000 gene set permutations and signal to noise ranking metric. We downloaded KEGG Pathways from the Molecular Signatures Database, which has 186 gene sets. After filtering for the recommended minimum (15) and maximum (500) gene set size, the remaining 145 gene sets were compared to the expression data from our dataset to calculate the GSEA enrichment score and to compute significant enrichment.

### Electrophysiology

Mice were anesthetized with chloral hydrate (400 mg/kg i.p.) and placed into a stereotaxic frame. Fur was plucked and an incision was made to expose the skull. Burr holes were drilled over the approximate location of the LC (from bregma, AP: 5.2-5.4, ML: 0.7-1.1).

Recordings were made using 16 channel silicone probes (A1×16-Poly2-5mm-50s-177-CM16LP, NeuroNexus) that were connected to a μ-series Cereplex headstage (Blackrock Microsystems). Digitized signals were acquired with a 16 channel Cereplex Direct system (Blackrock Microsystems) using a 250 Hz-5 kHz bandpass filter and a sampling rate of 10 kS/s. Probes were lowered to the approximate location of the LC (DV: −2.7-4.3).

LC units were identified based on standard criteria, including stereotaxic coordinates, biphasic response to foot-pinch/shock, and reduction/cessation of spontaneous activity following injection of the alpha-2 adrenergic receptor agonist clonidine (0.1 mg/kg, i.p.). For each set of recordings, a 5-min baseline period was collected, and was immediately followed by 10 applications of a contralateral foot-pinch separated by 10 s. Afterwards, 0.5 ms 1 mA footshocks were applied to the contralateral hindpaw separated by 10 s for 5.5 min to assess response to salient/aversive stimuli. LC spikes were manually sorted offline using Blackrock Offline Spike Sorting software.

Electrophysiology data was analyzed using Neuroexplorer. To ensure that recordings were from single units, neurons that had greater than 2% of recorded spikes within a predefined 3 ms refractory period were eliminated from the analysis. Basal firing rate and interspike intervals were calculated based on spikes collected within the 5-min baseline period. Spontaneous burst characteristics (number of bursts, percentage of spikes in a burst, burst duration, spikes per burst, interspike interval within a burst, burst rate, and interburst interval) during baseline was characterized using previously defined criteria derived from dopamine neurons (Grace and Bunney, 1983; Iro et al., 2021). Finally, response to footshock was analyzed in three time windows: immediate (0-60 ms), intermediate (60-100 ms), and long (200-400 ms), as previously described (Hirata and Aston-Jones, 1994).

### Behavioral assays

Behavioral assays were performed in the following order, from least to most stressful.

#### Novelty-induced and circadian locomotion

Individual mice were placed in a plexiglass arena (10” × 18” × 10”) surrounded by a 4 × 8 photobeam grid that records infrared beam breaks (Photobeam Activity System, San Diego Instruments, San Diego, CA; Lustberg et al. 2020). Two consecutive beam breaks were recorded as an ambulation, and total ambulatory activity was recorded. Mice were left undisturbed in the arena for 23 h. The first hour of the test reflected novelty responses while the remainder of the testing period showed changes in locomotion as a function of circadian cycle. The number and location of ambulations were recorded in 5-min intervals.

#### Novelty suppressed feeding

Chow was removed from individual home cages 24 h prior to behavioral testing. Mice were moved to the test room under red light and allowed to habituate for 2 h prior to the start of the test. Individual mice were placed in a novel arena (10” × 18” × 10”) with a single pellet of standard mouse chow located in the center. The latency to feed, operationally defined as grasping and biting the food pellet, was recorded using a stopwatch. Mice that did not feed within the 15-min period were assigned a latency score of 900s (Tillage et al., 2020).

#### Marble burying

Individual mice were placed in a novel arena (10” × 18” × 10”) containing 20 marbles of uniform size and color arranged in a 4 × 5 grid, each on top of 2” of lightly pressed cobb bedding. Mice were left undisturbed for 30 min in a brightly lit room. At the end of testing, the mice were placed back into home cages, and the number of marbles buried were counted by two independent observers. If different scores were reported between observes, the average was taken. A marble was considered buried if at least two-thirds of its height was submerged in the bedding. For each test cage, digital photographs were obtained at uniform angles and distances.

### Statistical analyses

Immunohistochemical quantification, stereological cell counting, and HPLC measurements of catechol concentrations were compared between saline and DSP-4 treated groups using a student’s t-test in GraphPad Prism. Similarly, behavioral assessment relied on t-test comparison between groups for total ambulations in the novelty-induced locomotion assay, latency to feed in the novelty-suppressed feeding task, and number of marbles buried in the marble-burying test. R Bioconductor packages were utilized for statistical analyses of RNA sequencing data, including differential gene expression (DGE). For electrophysiological recordings, students t-tests or Mann-Whitney tests were used for comparison of basal firing rates and spontaneous bursting properties between treatment conditions for normally and non-normally distributed data, respectively. A two-way repeated measures ANOVA, with response period as the within subject factor and treatment as the between subject factor, was used to analyze LC response to footshock.

## Results

### DSP-4 reduces NE content and dysregulates NE turnover in the pons and LC projection fields

To confirm the efficacy and specificity of DSP-4 (two injections of 50 mg/kg, administered one week apart), we assessed tissue levels of catecholamines and their metabolites (Fig. 1). DSP-4 dramatically reduced NE in the pons, where LC cell bodies reside (t_(6)_ = 17.73, p < 0.0001), as well as in the hippocampus (t_(14)_ = 3.27, p = 0.0056) and prefrontal cortex (PFC; t_(14)_ = 3.347, p = 0.0048), two of the primary projection regions of the LC. (Fig. 1a). DSP-4 treatment similarly decreased levels of MHPG, the primary catecholamine metabolite, in the pons (t_(6)_ = 5.205, p = 0.002), PFC (t_(14)_ = 2.978, p = 0.01) and hippocampus (t_(14)_ = 3.140, p = 0.0072) (Fig. 1b). Despite the reduction of NE, the rate of turnover (MHPG:NE ratio) was significantly increased in the pons (t_(6)_ = 2.685, p = 0.0363) and PFC (t_(14)_ = 2.499, p = 0.0255) compared to saline controls, with a similar trend seen in the hippocampus (t_(14)_ = 2.049, p = 0.0596), suggesting adaptations in the LC-NE system (Fig. 1c). By contrast, levels of other amine neuromodulators, including dopamine, serotonin, and their respective metabolites, were unchanged in all regions assessed (data not shown), confirming the specificity of this neurotoxin for noradrenergic neurons.

**Fig. 1.**
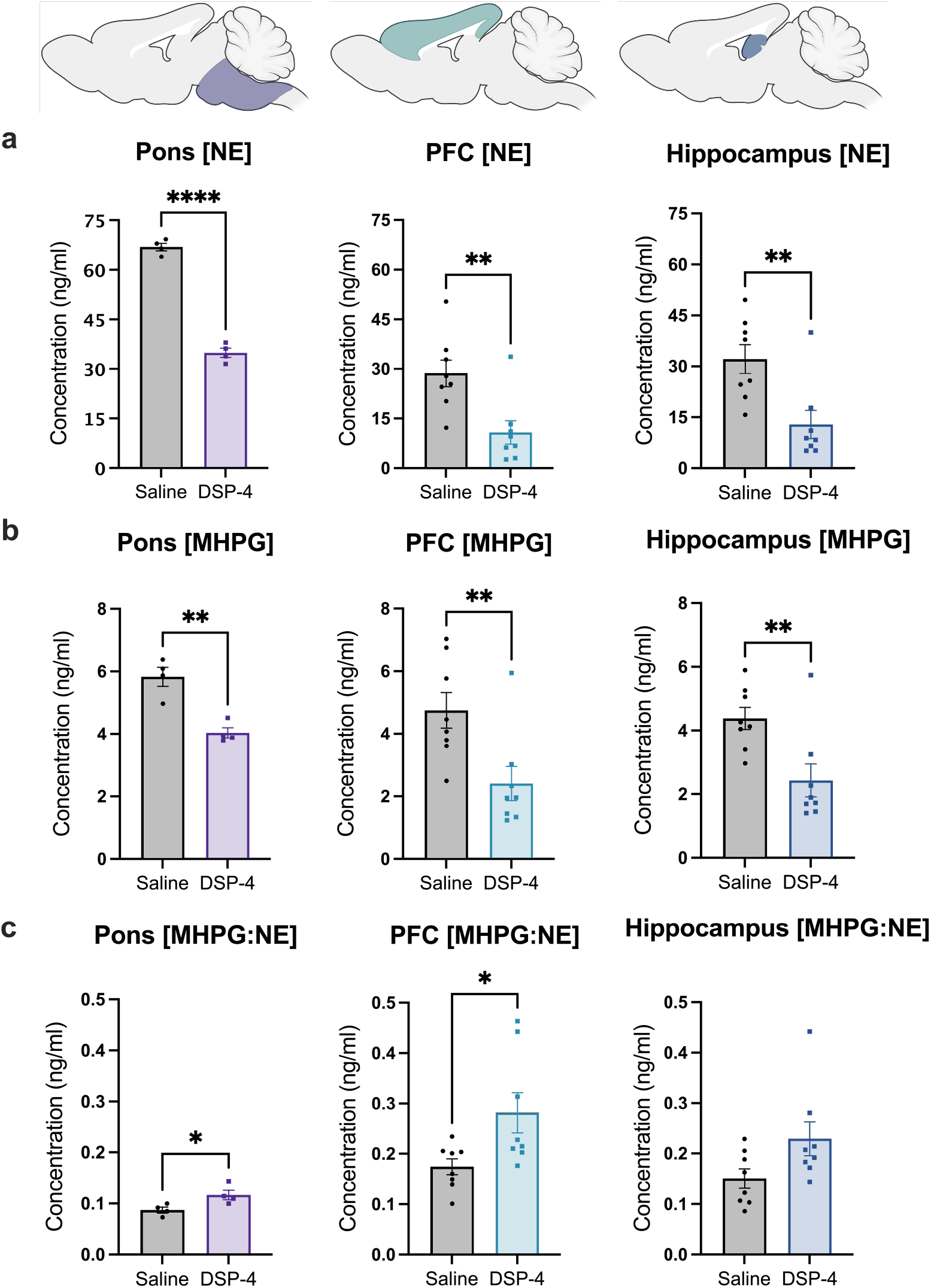
DSP-4 decreases tissue NE and metabolite levels and increases turnover. Mice received saline or DSP-4 (2 x 50 mg/kg, i.p.), and tissue monoamine and metabolite levels were measured 1 week later by HPLC in the pons, prefrontal cortex (PFC), and hippocampus (color shaded images at top represent the approximate regions dissected for analysis). DSP-4 significantly decreased NE (**a**) and its primary metabolite MHPG (**b**) in all 3 brain regions. (**c**) NE turnover, defined as the MHPG:NE ratio, was increased in the pons and PFC by DSP-4, with a similar trend in the hippocampus. Data shown as mean ± SEM. N=8 per group. *p<0.05, **p<0.01, ****p<0.0001.

### DSP-4 triggers loss of LC fibers and neuroinflammation but not frank cell body degeneration

Consistent with the depletion of tissue NE and canonical findings of LC fiber damage following DSP-4 treatment (Grzanna et al., 1989; Theron et al., 1993; Wolfman et al., 1994; Harro et al., 1999), NET immunoreactivity, a marker of LC axon, dendrite, and terminal integrity, was reduced in the LC (t_(6)_ = 6.003, p = 0.001), ACC (t_(6)_ = 13.76, p < 0.0001), and dentate gyrus (DG) region of the hippocampus (t_(6)_ = 10.23, p < 0.0001) in DSP-4 treated mice compared to controls (Fig. 2a). In contrast, we found that NET immunoreactivity in the bed nucleus of the stria terminalis (BNST), which receives noradrenergic innervation from brainstem A1 and A2 instead of the LC (Aston-Jones et al., 1999), was intact in DSP-4 treated mice (data not shown), indicating sparing of the ventral noradrenergic bundle and highlighting the specificity of DSP-4 induced damage to LC neurons.

**Fig. 2.**
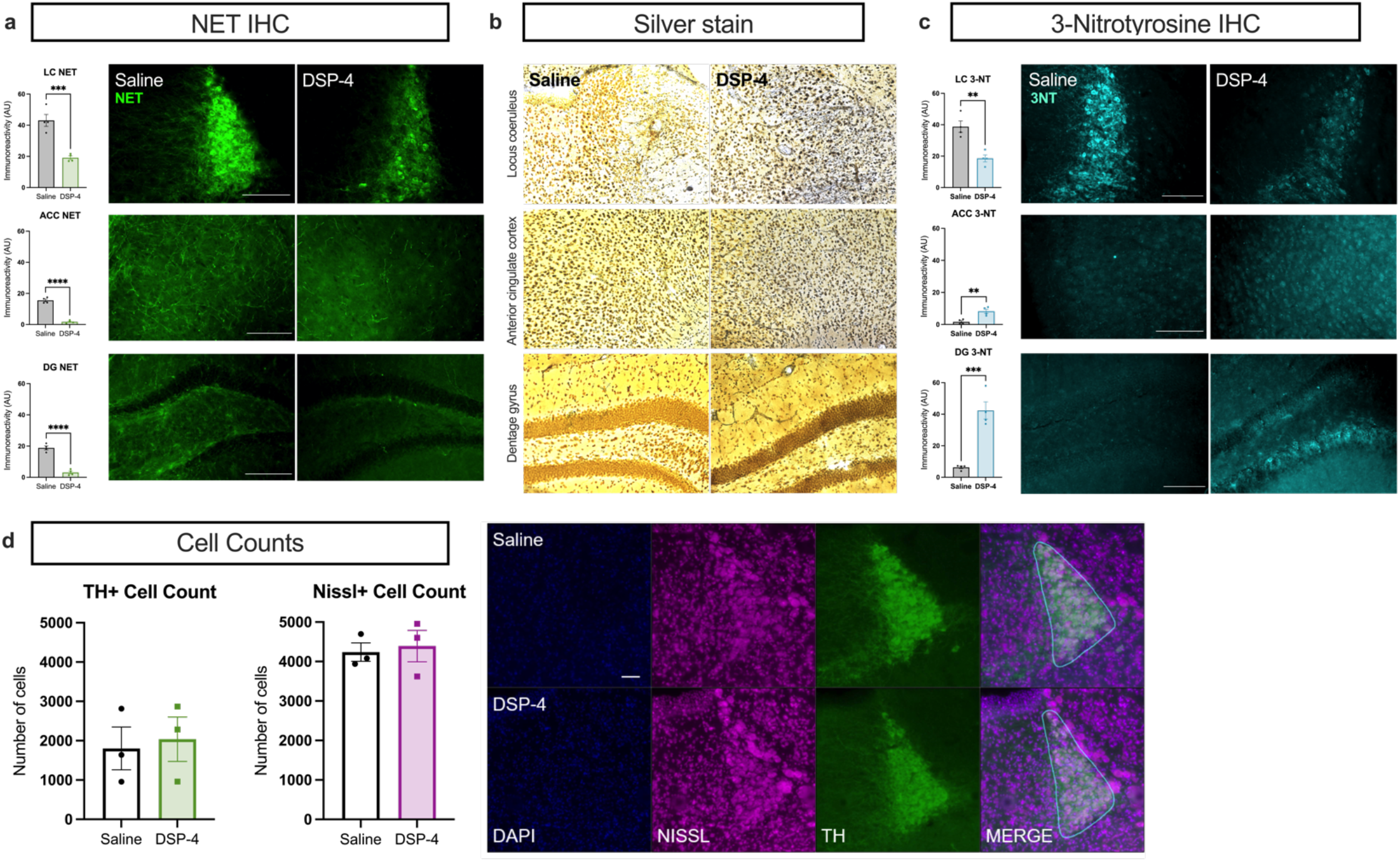
DSP-4 induces degeneration of noradrenergic terminals and oxidative stress but leaves LC cell bodies intact. Mice received saline or DSP-4 (2 x 50 mg/kg, i.p.) and assessed for locus coeruleus (LC) neuron damage 1 week later. (**a**) DSP-4 results in substantial loss of axon terminals as measured by norepinephrine transporter (NET) immunoreactivity in the dentate gyrus (DG) and anterior cingulate cortex (ACC), with a similar decrease in NET also present in the LC itself. (**b**) Representative images of silver stained brain tissue indicates neurodegenerative processes in the LC, ACC, and DG following DSP-4. (**c**) The oxidative stress marker 3-nitrotyrosine (3-NT) was increased in the ACC and DG but decreased in the LC by DSP-4. (**d**) Despite NE fiber damage and the evidence of neurodegenerative and oxidative processes following DSP-4, LC cell body number was unaffected as measured by TH and NeuroTrace Nissl immunoreactivity. Data shown as mean ± SEM. N=3-4 per group. **p<0.01, ***p<0.001, ****p<0.0001.

Next, we used silver staining to verify that the loss of NET immunoreactivity reflected LC fiber degeneration and not just a downregulation of NET. Robust silver staining in the LC, ACC, and DG indicated the presence of degenerative processes in the LC and projection regions (Fig. 2b). Finally, we assessed levels of oxidative stress marker 3-NT in the LC and projection regions and found that it was significantly decreased in the LC following DSP-4 administration (t_(6)_ = 4.746, p = 0.0032), but was increased in the ACC (t_(6)_ = 4.397, p = 0.0046) and DG (t_(6)_ = 6.558, p = 0.0006) (Fig. 2c).

To determine whether damage resulting from DSP-4 impacted cell body integrity, we performed a count of LC neurons. We quantified DAPI+ cells also positive for TH or NeuroTrace Nissl in the LC. We found no differences in the number of LC neurons in DSP-4 treated mice compared to controls in either analysis (Fig. 2d).

Because neuroinflammation often occurs in response to damage and is a key component of AD and PD pathology (Tansey et al., 2022; Thakur et al., 2022), and glial activation can be impeded by NE signaling (Liu et al. 2019), we assessed immunoreactivity of the astrocyte marker GFAP and the microglial marker Iba-1 in the LC and target regions. We observed robust astrocytic (t_(4)_ = 6.969, p = 0.0022) and microglial (t_(4)_ = 3.642, p = 0.0219) responses in and around the LC of DSP-4 treated mice compared to controls (Fig. 3). GFAP immunoreactivity was elevated in the DG (t_(6)_ = 2.227, p = 0.0675), but decreased in the ACC (t_(6)_ = 2.480, p = 0.0478), and Iba-1 was increased in both the ACC (t_(6)_ = 3.351, p = 0.0154) and the DG (t_(6)_ = 4.118, p = 0.0062). These results reveal that damage to the LC and its projections recapitulates key aspects of neuroinflammation and neurodegeneration in AD and PD.

**Fig. 3:**
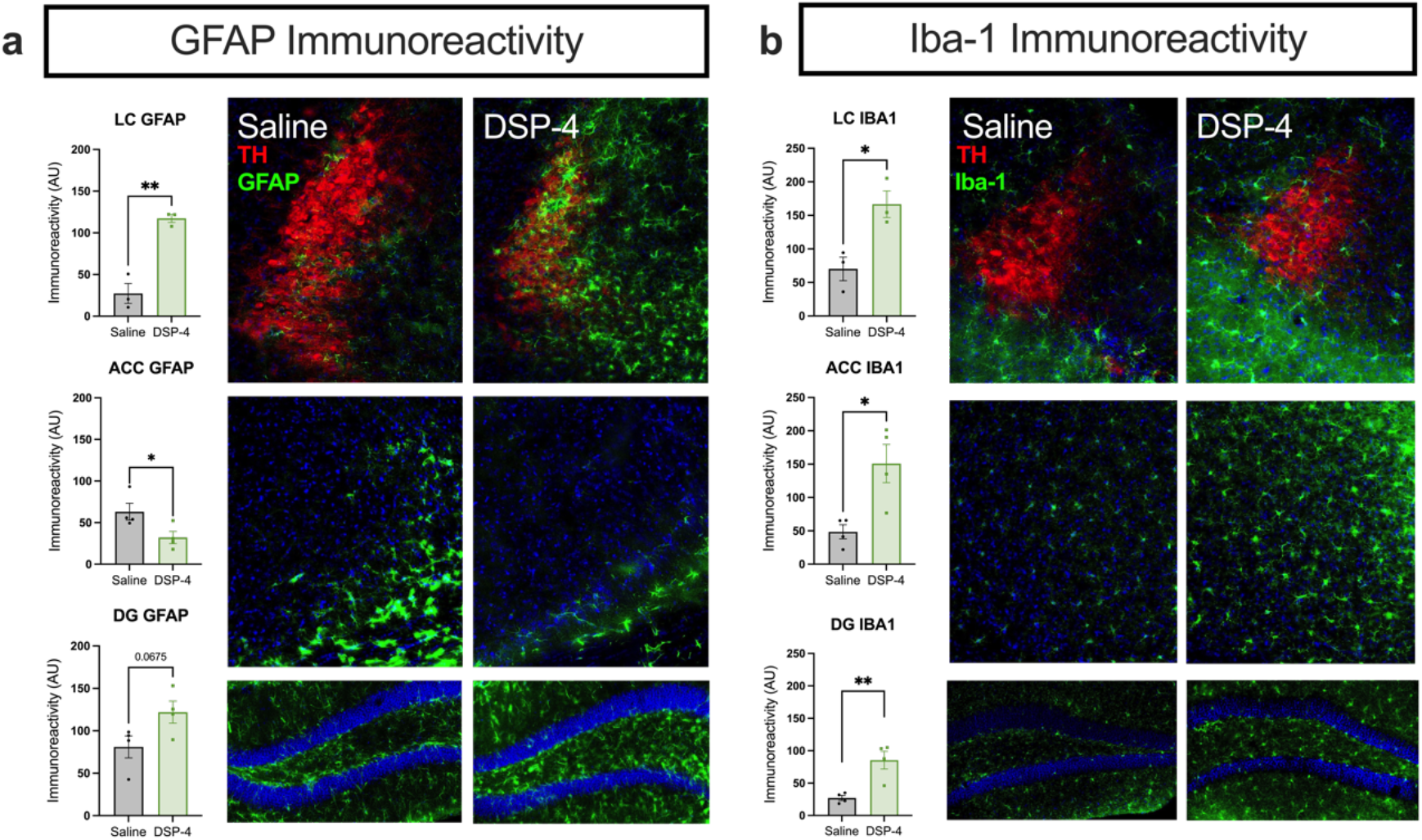
DSP-4 alters astrocyte and microglia activation across the LC-NE system. Mice received saline or DSP-4 (2 x 50 mg/kg, i.p.) and assessed for neuroinflammation 1 week later. (**a**) Astrocyte reactivity, as measured by GFAP immunostaining, was significantly increased in the locus coeruleus (LC) with a trend in the dentate gyrus (DG) following DSP-4 treatment, while the anterior cingulate cortex (ACC) showed a decreased astrocytic response. (**b**) Microglial response, indicated by Iba-1 immunoreactivity, was increased across all regions assessed. Data shown as mean ± SEM. N=3-4 per group. *p<0.05, **p<0.01.

### DSP-4 treatment leads to molecular but not cellular dysfunction in LC cell bodies

The marked reduction in NET immunoreactivity (Fig. 2a) in the LC without the loss of cell bodies (Fig. 2d) suggested that dysregulation of noradrenergic markers may be occurring first on a molecular level. To assess this, we expanded our analysis to the entire LC transcriptome as a complement to our immunohistochemical staining of specific protein markers. We employed a *Slc2a6-Rpll10-eGFP* line for specific targeting of noradrenergic neurons through TRAP (Mulvey et al., 2018). We observed robust enrichment of LC genes in our TRAP samples compared to input (Fig. 4a and data not shown), indicating the successful implementation of this technique. Next, we assessed differentially expressed genes (DEGs) between replicate samples from the DSP-4 treatment and saline control groups. Most notably, we saw a marked downregulation of multiple noradrenergic function and specification genes, including *Slc6a2* (NET; logFC = −1.5032, p < 0.0001), *Th* (logFC = −0.7850, p = 0.0006), *Dbh* (logFC = −1.2642, p < 0.0001), and *Phox2a* (logFC = −1.0292, p = 0.0054), and the LC-enriched neuropeptide *Gal* (logFC = −1.1636, p < 0.0001), which is reflective of a loss of LC neuron “identity” following DSP-4 administration (Fig. 4b and 4c). While only three DEGs reached stringent statistical significance with a false discovery rate (FDR) < 0.1 *(Slc6a2, Dbh, Gal),* this experiment yielded a substantial list of biologically informative transcripts (Figure 4c). Further analysis of gene expression networks using WGCNA revealed four modules that were significantly correlated with treatment, one of which included *Dbh, Gal,* and *Slc6a2* (Fig 4d). This module contained 72 genes, including several that are implicated in neurodegenerative diseases, suggesting that critical LC genes (*Dbh*, *Gal*, *Slc6a2*) are clustering with neurodegeneration genes in their expression patterns after treatment with DSP-4. Finally, using GSEA to compare the gene expression patterns in our dataset with repositories of known gene sets, we identified 17 KEGG pathways that were significantly enriched in our dataset, including oxidative phosphorylation (enrichment score (ES)=-0.53, p<0.001), lysosome (ES=0.41, p=0.006), pathways in cancer (ES=0.35, p=0.004), melanogenesis (ES=0.44, p=0.023), and cytokine-cytokine receptor interaction (ES=0.45, p=0.025). Notably, the pathways for PD (ES=-0.49, p<0.001), AD (ES=-0.37, p=0.037), and Huntington’s disease (ES=-0.45, p=0.002) were significantly negatively correlated with treatment, and further investigation revealed that many of the core enrichment genes in these pathways are similarly downregulated in clinical neurodegenerative diseases and after DSP-4 treatment in mice (Fig. 4e).

**Fig. 4.**
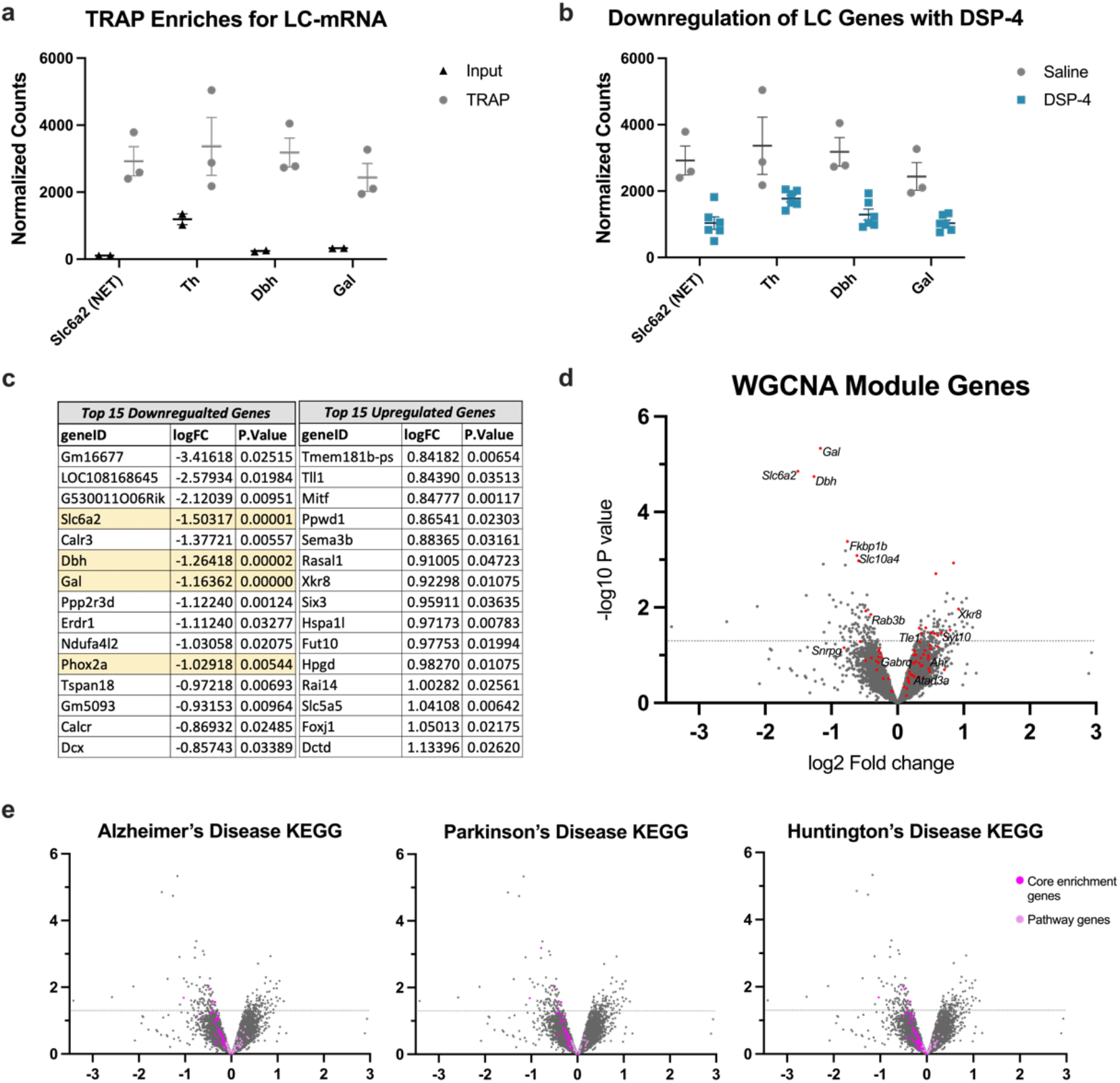
DSP-4 triggers changes in the LC transcriptome. Mice received saline or DSP-4 (2 x 50 mg/kg, i.p.) and LC gene expression was assessed 1 week later. (**a**) TRAP allows for purification of mRNA from LC neurons through immunoprecipitation (TRAP, grey), resulting in enrichment of noradrenergic genes compared with mRNA from the entire hindbrain sample (input, black). (**b**) Differential gene expression (DGE) was assessed between DSP-4 (blue) and saline control (grey) TRAP samples, and revealed that noradrenergic-specific genes, including galanin (*Gal*), norepinephrine transporter (*Slc6a2*), dopamine ß-hydroxylase *(Dbh)* and tyrosine hydroxylase (*Th*), were among the most significantly and robustly downregulated transcripts in the LC of DSP-4 treated mice. Data for **a** and **b** shown as mean ± SEM, N=2-6 per group. (**c**) List of top 15 downregulated and top 15 upregulated DGE, sorted by fold change (logFC) with p < 0.05. (**d**) Volcano plot of all filtered, normalized genes (~11,500) with genes from WGCNA-defined module in red. Labeled genes from this module are those of interest based on published connections to neurodegenerative disease. (**e**) Volcano plots (as shown in (d)) highlighting genes from 3 significantly enriched KEGG pathways in our LC data, with GSEA-identified “core enrichment genes” colored magenta, and remaining pathway genes colored light pink.

Next, we investigated whether these molecular changes in LC neurons after DSP-4 treatment were accompanied by cellular changes. LC neurons are tonically active, show elevated tonic activity during stress, and exhibit “bursting” activity in response to salient and novel stimuli (Valentino, 1988; Vankov et al., 1995; Curtis et al., 1997; McCall et al., 2015). To determine whether DSP-4-induced molecular dysregulation impacted cellular activity, *in vivo* electrophysiology under anesthesia was conducted to measure baseline and footshock-evoked firing of LC neurons. There were no differences in the baseline firing rate of LC neurons between treatments (Mann-Whtiney U = 1072, p = 0.2312) (Fig. 5), with interspike intervals and spontaneous bursting properties also remaining unchanged (data not shown). There was a main effect of time period in response to footshock such that firing rate of LC neurons decreased in each successive response phase (F_2,176_ = 17.09, p < 0.0001). However, there was no main effect of treatment (F_1,188_ = 2.408, p = 0.1243) or a treatment x time period interaction (F_2,176_ = 0.2655, p = 0.7671) on response to footshock. We conclude that the molecular changes occurring following DSP-4 do not significantly affect LC neuron firing under these conditions.

**Fig. 5.**
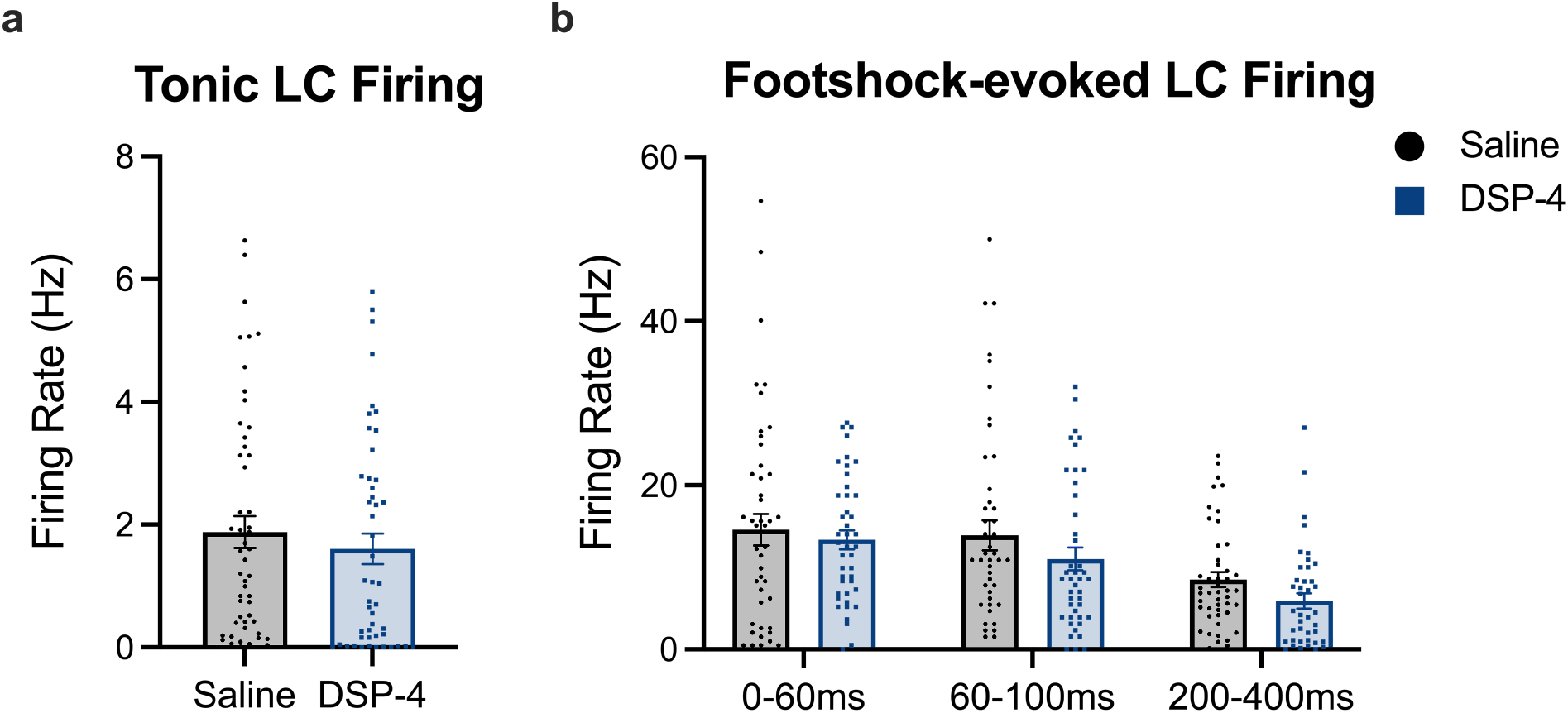
DSP-4 treatment does not alter baseline or footshock-induced LC activity. Mice received saline or DSP-4 (2 x 50 mg/kg, i.p.) and assessed for locus coeruleus (LC) neuron firing under anesthesia 1 week later. No differences were found for (**a**) baseline tonic firing rates (in Hz). (**b**) There was no main effect of treatment on footshock-evoked firing rates of LC neurons 0-60, 60-100, or 200400 ms following the stimulus. Data shown as mean ± SEM. N=5 mice per group 42-53 neurons/group. *p<0.05

### DSP-4 treatment results in a novelty-induced anxiety phenotype, implying compensatory hyperactivity of LC-NE transmission

AD and PD share several prodromal behavioral symptoms related to affect and arousal, processes known to be regulated by the LC-NE system and sensitive to LC integrity (Weinshenker, 2018). Moreover, cognitive impairment is a diagnostic criterion for AD and is common in later stage PD; thus, we assessed the consequences of DSP-4 on LC/NE-sensitive behaviors that reflect prodromal and cognitive abnormalities in AD and PD. We found that lesioned mice were profoundly more reactive in novelty-induced stress paradigms, which are commonly used to model anxiety and are bidirectionally modulated by NE (Lustberg et al., 2020b). DSP-4 treated mice took significantly longer to consume food in the novelty-suppressed feeding test (t_(20)_ = 3.158, p = 0.0048), buried more marbles (t_(14)_ = 4.290, p = 0.0007), and ambulated less during the first hour in a novel cage (t_(14)_ = 2.999, p = 0.0096) compared to saline-treated controls (Fig. 6). Importantly, DSP-4 treatment had no effect on latency to eat in the home cage or in total ambulations across a 23-h period, suggesting increased anxiety-like behavior rather than a decrease in hunger or general locomotion. Arousal (as assessed by latency to fall asleep following gentle handling) and associative memory (as measured by freezing in a footshock-associated context) did not differ between treatment groups (data not shown).

**Fig. 6.**
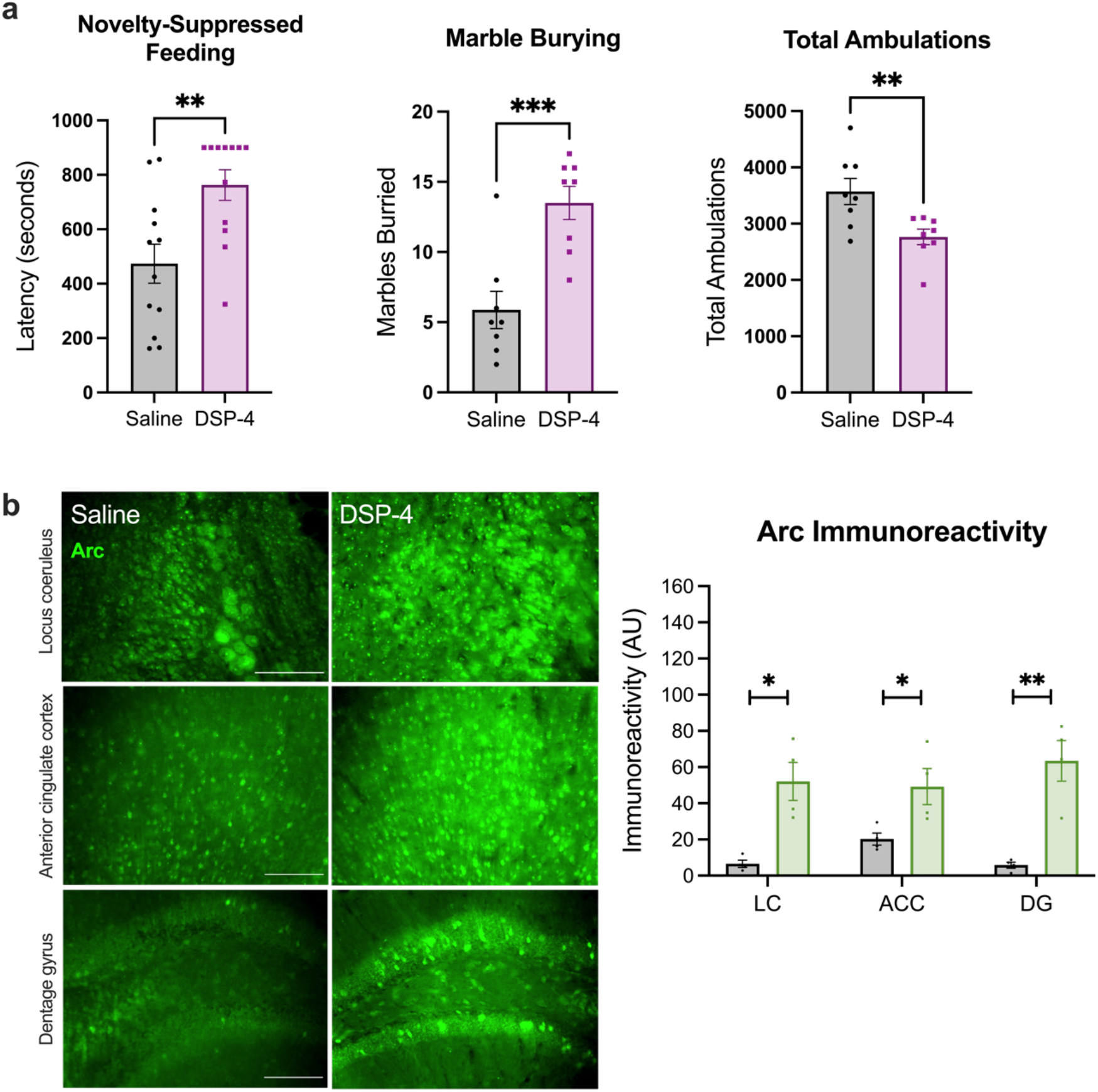
DSP-4 increases novelty-induced anxiety and arc expression. (**a**) DSP-4-treated mice display increased latency to bite the food pellet in the novelty-suppressed feeding test, buried more marbles, and showed fewer ambulations in response to a novel environment (N=8-12 per group). (**b**) Immunoreactivity for the immediate early gene Arc following cage change was reduced in the locus coeruleus (LC) and projection regions (anterior cingulate cortex, ACC; dentate gyrus region of the hippocampus, DG) (N=3-4 per group). Data shown as mean ± SEM. *p<0.05, **p<0.01, ***p<0.001.

Elevated novelty-induced anxiety-like behavior is consistent with increased noradrenergic activity (Lustberg et al., 2020a), which was surprising given the profound loss of noradrenergic fibers and NE in LC terminal fields. Our HPLC data indicated increased NE turnover from surviving terminals, and adrenergic receptor supersensitivity has been reported in DSP-4 lesioned animals (Wolfman et al., 1994; Szot et al., 2010). To investigate whether these adaptations are sufficient to boost downstream signaling mechanisms that underlie behavioral reactivity, immunostaining for Arc was quantified following cage change, a mild stressor that is resistant to habituation and sensitive to NE transmission (Vankov et al., 1995; McCall et al., 2015; Takeuchi et al., 2016; Grella et al., 2019; Lustberg et al., 2020b; Lustberg et al., 2020a; Prokopiou et al., 2022). Arc is an immediate early gene and neural activity marker that is induced following adrenergic receptor stimulation, and acts as a readout of postsynaptic LC-NE transmission (McIntyre et al., 2005; McReynolds et al., 2014). After cage change, a marked increase in Arc immunoreactivity was observed in the LC (t_(6)_ = 4.258, p = 0.0053), ACC (t_(6)_ = 2.755, p = 0.0331), and DG (t_(6)_ = 5.099, p = 0.0022) of DSP-4 treated mice compared to saline-treated mice (Fig. 6).

## Discussion

The present study characterized the impact of DSP-4 on the molecular, cellular, and behavioral levels in mice to comprehensively assess the consequences of damage to LC neurons reminiscent of early stages of AD and PD. The depletion of NE and its metabolite MHPG, as well as the elevated MHPG:NE ratio, by DSP-4 are consistent with decades of previous research and indicate reduced total NE but increased NE turnover (Jonsson et al., 1981; Logue et al., 1985; Harro et al., 1999; Szot et al., 2010). Immunohistochemical staining following DSP-4 treatment revealed a dramatic reduction of NET in the PFC and hippocampus, which could mean a loss of noradrenergic terminals and/or a downregulation of NET expression. We found evidence for both. NET mRNA in LC neurons was significantly reduced by DSP-4, and silver staining provided evidence for degeneration of axons/terminals in projection regions. Diminished NET immunoreactivity was also observed in the LC of DSP-4 treated mice. To determine whether this was due to a loss of neurons, we counted LC cell bodies (i.e. positive for DAPI and TH or Nissl) and found no effect of DSP-4, which is typical for similar dosing regimens (Lyons et al., 1989; Matsukawa et al., 2003; Szot et al., 2010). Taken together, the loss of NE and NET in terminal regions and pons is indicative of LC axon, terminal, and dendrite degeneration, with the increased MHPG:NE ratio potentially signifying compensatory elevation of NE release from surviving fibers (Jacobs, 2019; van Hooren et al., 2021; Gilvesy et al., 2022).

DSP-4 provoked oxidative stress in the forebrain and a robust neuroimmune response in the both the LC and its projection regions. 3-NT immunoreactivity was elevated in the ACC and DG following DSP-4 administration, providing a link to AD, as LC lesions also increased 3-NT in the cortex of mice that overexpress mutant amyloid precursor protein (Heneka et al., 2006). Paradoxically, the abundance of 3-NT in control LCs was high at baseline relative to the other brain regions and decreased by DSP-4. Catecholamine synthesis and metabolism generate oxidative stress, which could contribute to the high baseline levels of 3-NT in the LC, while the DSP-4 induced reduction may reflect the loss of catecholamine synthetic capacity. Interestingly, lipopolysaccharide-induced 3-NT oxidative stress in the dopaminergic SN that is reminiscent of PD pathology was also attenuated by DSP-4 lesions of the LC (Iravani et al., 2014).

Microglial activation as measured by Iba-1 immunoreactivity was elevated in the LC, ACC, and DG of DSP-4 treated mice compared to controls. GFAP+ reactive astrocytes were increased in the LC and DG of DSP-4 tissue but were suppressed in the ACC. The interplay between neuroinflammation and neurodegeneration is bidirectional, and our experiments with NeuroSilver staining revealed active degenerative processes in both fibers and cell bodies of the LC and its projection regions. Because our analysis was restricted to a single timepoint when both processes were evident, we cannot know whether one triggered the other. Neuroinflammation and neurodegeneration are key components of AD and PD, and both are exacerbated by ablation of LC-NE in animal models of these disorders and can be ameliorated by pro-noradrenergic therapies in clinical populations (Rommelfanger et al., 2004; Heneka et al., 2006; Rommelfanger et al., 2007; Heneka et al., 2010; Chalermpalanupap et al., 2018; Song et al., 2019a; Song et al., 2019b; Levey et al., 2022). Given that we observed no difference in LC cell body number, our DSP-4 dosing regimen appears to represent an early phase of neurodegeneration, which is characterized by initial loss of noradrenergic innervation prior to frank LC degeneration in AD and PD (Fritschy et al., 1990; Doppler et al., 2021; Gilvesy et al., 2022).

In AD and PD, LC neurons become dysfunctional early on but persist for many years prior to cell death. However, almost nothing is known about the molecular changes that drive LC dysfunction prior to outright degeneration. Due to its small size and neuron number, selective mRNA profiling of the murine LC transcriptome has historically been challenging. To address this gap, we assessed the effects of DSP-4 treatment on the LC transcriptome using TRAP and obtained enrichment of known LC genes such as *Th, Slc6a2* (NET), *Dbh, Phox2a,* and *Gal* in saline-treated mice, as previously reported (Mulvey et al., 2018). Remarkably, we found that these same genes were down-regulated in DSP-4 treated mice, suggesting a deterioration of noradrenergic identity prior to LC neuron loss. Differential gene expression and down-regulation of NET, *Dbh,* and *Th* are consistent with *in vitro* studies exposing SH-SY5Y cells to DSP-4 (Wang et al., 2014), but opposite of what has been reported in clinical AD, where *Th* and NET are increased in surviving LC neurons (Szot et al., 2006). We speculate that compensatory increases in noradrenergic markers are triggered by the cell loss ubiquitous in late-stage AD, which did not occur in our DSP-4 treat mice. In addition to the disruption in noradrenergic gene expression, we also found several genes associated with neurodegeneration and neurotoxicity that were differentially expressed between our treatment groups. Some of these genes, highlighted in Fig. 4 and detailed in Table 2, were also found in the co-expression network module with *Gal*, *Dbh*, and *Slc6a2*. These results were complemented by significant enrichment in gene sets associated with AD, PD, and Huntington’s disease, and contribute to the rich array of potential genetic targets for better understanding the early stages of neurodegeneration.

**Table 2.**
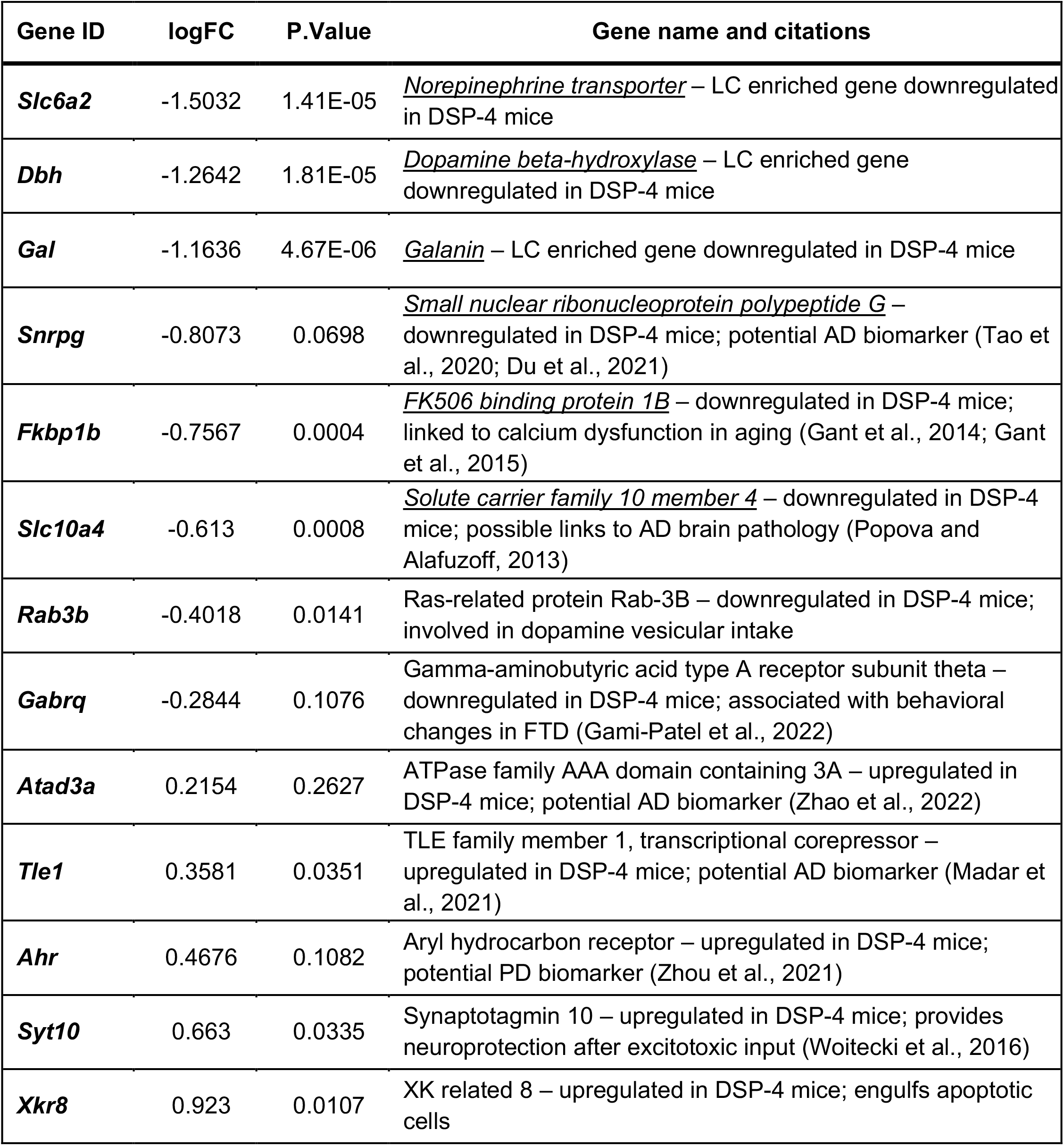
Highlighted genes from WGCNA shown in Figure 4.

Prodromal AD and PD are characterized by neuropsychiatric abnormalities including anxiety, agitation, depression, and sleep disturbances that appear long before the primary diagnostic symptoms of these diseases (cognitive and motor impairment, respectively). Using a battery of tests to probe these behavioral domains, we found that DSP-4-treated mice displayed increased anxiety-like behavior specific to novelty: they took significantly longer to consume a food pellet, buried more marbles, and had reduced locomotor activity in novel environments. These results are consistent with previous reports that surgical or neurotoxin ablation of the LC in rats induces similar anxiety-like phenotypes that reflect responses to novelty stress (Martin-Iverson et al., 1982; Harro et al., 1995).

LC-NE transmission is triggered by novelty stress, and activation of this system promotes, while inactivation suppresses, novelty-induced anxiety (McCall et al., 2015; Lustberg et al., 2020b; Lustberg et al., 2020a). Thus, the emergence of anxiety-like behavior in DSP-4-treated mice and AD/PD patients where a dramatic *loss* of noradrenergic fibers and NE is evident creates a paradox and suggests that compensatory mechanisms are engaged in response to LC damage that lead to hyperactive NE transmission. We can imagine three potential neuroanatomical/neurobiological substrates where this compensation may occur: LC cell bodies (e.g. increased neuron firing), LC terminals (e.g. increased NE release), and/or postsynaptic compartments (e.g. receptor/signaling molecules super-sensitivity).

Using in vivo electrophysiology, we detected no differences in the baseline (firing rate, interspike interval, spontaneous bursting properties) or footshock-evoked firing rate of LC neurons. This is consistent with previous reports in DSP-4 treated rats (Szot et al., 2010) but distinct from partial 6-OHDA LC lesions, which increased LC activity in mice (Szot et al., 2016). One important difference is that the 6-OHDA-treated mice had LC neuron loss (~30%), while LC cell bodies remained intact in our study. These results suggest that LC neurons are capable of compensatory increases in firing, but that cell body degeneration is required to trigger this response. By contrast, we did find evidence for increased NE release. Although we did not measure this directly, metabolite to parent neurotransmitter ratio is a validated proxy for turnover. We detected elevated MHPG:NE ratio in the LC and terminal regions, which has been previously reported in DSP-4 treated rodents (Hallman and Jonsson, 1984) and consistent with human AD cerebrospinal fluid data (Francis et al., 1985; Hoogendijk et al., 1999; Raskind et al., 1999; Jacobs et al., 2021b). Indeed, high CSF MHPG levels are associated with neuropsychiatric abnormalities in AD (Jacobs et al., 2021a).

Finally, postsynaptic compensatory mechanisms resulting from the loss of LC fibers may be at play. There are many reports of increased adrenergic receptor density following DSP-4 administration (Johnson et al., 1987; Harro et al., 1999), but the consequences on downstream receptor signaling have not been carefully investigated. We performed immunostaining for the immediate early gene Arc, which is a marker for neuronal activity and induced by activation of adrenergic receptors (Essali and Sanders, 2016). Following cage-change stress, Arc immunoreactivity was dramatically elevated in DSP-4 treated tissue compared to controls in the LC and its output regions in the forebrain. We conclude that compensatory changes in NE release from surviving LC terminals and/or postsynaptic adrenergic receptor signaling could contribute the anxiogenic effects of DSP-4, while increases in LC firing do not. These results have important implications for AD and PD, where early LC pathology, damage to noradrenergic fibers, and neuropsychiatric symptoms precede frank LC loss in prodromal disease. In addition, these experiments support the notion that DSP-4 dysregulates, rather than simply ablates, the noradrenergic system, and should act as a caution to researchers employing this neurotoxin as they interpret their results.

## Acknowledgements

This work was supported by a National Institutes of Health (NIH) National Institute of Environmental Health Sciences Training Grant (ES12870) and an NIH D-SPAN F99/K00 Award from the National Institute of Neurological Disorders and Stroke (NS129168) to A.F.I, as well as RF1 awards from the National Institute on Aging (AG079199 and AG061175) to D.W. This study was supported in part by the Emory HPLC Bioanalytical Core (EHBC), which is subsidized by the Emory University School of Medicine and is one of the Emory Integrated Core Facilities. Additional support was provided by the Georgia Clinical & Translational Science Alliance of the National Institutes of Health under Award Number UL1TR002378. We thank Q. Eastman for helpful manuscript editing and Dr. M. McGuirk Sampson for assistance with R code for analysis.

## Contributions

A.F.I. designed and performed research, analyzed data, and wrote the manuscript. M.A.K. helped design and performed research and analyzed data and helped write and edit the manuscript. D. L. helped design and performed research and analyzed data. A.K. performed research and analyzed data. K.E.M. analyzed data and assisted with writing and editing portions of the manuscript. B.M. designed research and analyzed data. A.S. analyzed data. L.C.L. assisted with research design and analysis. S.A.S. assisted with research design and data analysis. J.D.D. assisted with research design and contributed analytic tools. D.W. designed research and wrote the manuscript.

